# Changes of TLQP-21 in Response to Glucose Load

**DOI:** 10.1101/626283

**Authors:** Giulia Corda, Barbara Noli, Barbara Manconi, Carla Brancia, Manuela Pellegrini, Fabio Naro, Gian-Luca Ferri, Cristina Cocco

**Affiliations:** NEF-Laboratory, Department of Biomedical Sciences, University of Cagliari, Monserrato (CA), Italy; Department of Life and Enviromental Sciences, University of Cagliari, Monserrato (CA), Italy; Department of Anatomical, Istological and Legal medicine Sciences of the locomotor apparatus, University of “La Sapienza’, Roma, Italy

**Keywords:** TLQP-21, glucose, insulin, VGF, metabolism

## Abstract

The TLQP-21 peptide peripherally potentiates glucose-stimulated insulin secretion. The aim of this study was to investigate a possible endocrine mechanism through which TLQP-21 increases the insulin secretion. Using an antibody specific for the common N-terminal portion of the TLQP peptides, we studied pancreas and plasma of mice subjected to intraperitoneal glucose load, by immunohistochemistry and immunosorbent assay (ELISA), alone or coupled to High Performance Liquid Chromatography (HLPC). Mice underwent a period of starvation hence have received a glucose load, or saline, and were sacrificed 30 or 120 minutes later. In normal endocrine pancreas, the TLQP-antiserum stained either peripheral or central cells. Interestingly, 30 min after a glucose load, TLQP immunostaining was disappeared in pancreas and, when analysed by ELISA, the TLQP-levels started to increase in plasma reaching peak concentration at 120 min. (controls *vs.* 30 and 120 min.: *p*<0.05 and *p*<0.001, respectively). The analysis of plasma and pancreas extracts using HLPC coupled to ELISA demonstrated the presence of the TLQP-21, with statistically significant increase of this peptide in plasma at 120 min (*vs*. controls p<0.05) in agreement to the changes seen by measuring the totality of the TLQP peptides. In pancreas sections we found the presence of the C3a-R1, involved in insulin secretion, and previously identified as a TLQP-21 receptor. Hence, after a glucose load, TLQP-21, may be released by the pancreas into the plasma, returning to the pancreas in order to modulate the insulin secretion through the C3a-R1.

## INTRODUCTION

Insulin secretion includes a pattern of paracrine, autocrine, endocrine, and autonomic neural signaling mechanisms. The VGF precursor/proprotein (617 or 615 amino acids in rats/mice and humans, respectively) [1], codified from *vgf* gene (no acronym name), contains a large amount prohormone convertases cleavage sites PC1/3 and PC2 [2], from which a number of peptides of different molecular weight (MW) are derived [3], including the VGF derived peptide TLQP-21 (VGF_556-564_). The C-terminal internal fragment TLQP-21 was initially studied for its role in energy balance [4 – 7]. The TLQP-21 is expressed in the pancreas [8, 9] and it is able to potentiate the glucose stimulated insulin secretion, in both human and rat pancreatic islets [10, 11]. When peripherally injected into rats, this peptide is able to reduce the blood-glucose peak (after about 20 min. from the bolus ingestion) and to increase the plasma insulin levels. Moreover, it preserves the islet mass and slows the onset of diabetes in Zucker Diabetic Fatty rats. Overexpression of VGF in primary rat islets resulted in a 46% increase in glucose-stimulated insulin secretion, without affecting basal insulin secretion [11]. The TLQP-21 action mechanisms are not clarified yet, but two different G-coupled protein receptors have been discovered as TLQP-21 receptors. The complement component 3a receptor (C3a-R1) [12 – 14] is involved with the TLQP-21 in modulating the lipolysis [14], while the receptor for the globular heads of C1q (gC1q-R) acts together with the TLQP-21 as a modulator of neuropathic pain [15]. The modality of action with which TLQP-21 acts in potentiating insulin secretion has not been established yet. Hence, the aim of the present study was to investigate the possible endocrine activity of TLQP-21 in response to glucose stimuli. So, we analyzed possible alterations of the TLQP-21 in both plasma and endocrine pancreas through immunohistochemistry (IHC) and immunosorbent assay (ELISA) alone or coupled to High Performance Liquid Chromatography (HLPC).

## MATERIAL & METHODS

### Animals and tissue samples

Male CD1 mice (Charles River, Lecco, Italy; aged 12 weeks, weight 35–45 g) have been housed (6 animals/cage) at the enclosure of the University of Cagliari under controlled temperature (23 ±2°C), light (12-12h; light phase: from 8am to 8pm and dark phase: from 8pm to 8am) and relative humidity (60 ±10 %) conditions. Mice underwent a period of starvation 18h before the glucose tolerance test and the sacrifice. The animals have been kept with free access to water. Mice (n=12 per group) have been injected with glucose (3g/kg) or saline and sacrificed 30 or 120 min. after. Immediately before the sacrifice, a peripheral venous blood sample (from tail vein) was obtained for the measurement of blood glucose level (mg/ml, MultiCare-in, Biochemical System, S.r.l, Arezzo, Italy). For the sacrifice, mice have been anesthetized (under gas anesthesia induced by isofluran, IsoFlo, Italy) and blood (100-200μl) has been drawn from the left heart with syringes pretreated with ethylenediaminetetraacetic acid disodium salt (EDTA; 1.78 mg/ml). The blood samples collected have been drawn into tubes, centrifuged (11,000 rpm; 10 min.) and frozen until use. After blood collection, the perfusion has been performed with 40 g/L paraformaldehyde (40 g/L in 0.1 mol/L PO_4_, 15 min.) injected into the left ventricle. Fixed pancreas samples have been extracted from the body and rinsed in phosphate buffer saline (PBS: 0.01 mol/L PO_4_, pH 7.2, 0.15 mol/L NaCl) containing 70g/L sucrose and 0.1g/L NaN_3_, treated with cryoembedding media (PVA 56–98 59 g/L, Tween-20 10 g/L, and Peg-400 40 g/L in PBS-NaN_3_ 1 g/L)[16] and subsequently frozen with liquid nitrogen. Cryosections (10 μm) obtained with cryomicrotome (Microm HM-560, Walldof, Germany) [17] have been collected on slides coated with poly-L-lysine (Sigma, Milan, Italy) and stored until use. Additional control rats were used for HPLC coupled to ELISA (n=6), sacrificed, under gas anesthesia with isofluran, by decapitation. Pancreas samples were obtained and extracted in PBS containing protease inhibitor cocktail (P8340, Sigma-Aldrich, Schnelldorf, Germany), homogenized using an Ultra-Turrax Homogenizer (Ika-Werke, Staufen, Germany), heated in a boiling water bath (10-15 min), centrifuged (3,000 rpm, 10-15 min) and frozen until use. Experimental protocols were approved by the Ethical Committee of the University of Cagliari and were performed in agreement with the Italian legislation, while the care and use of animals were approved by the American Physiological Society and EEC council directive of 24 November 1986 (86/609). All surgery was performed under sodium pentobarbital anesthesia, and all efforts were made to minimize suffering.

### TLQP antiserum

The guinea-pig primary antiserum against TLQP peptides specific for their common N-terminal portion, previously described in detail [18] was extensively used in different rat organs and tissues [8, 19 – 21]. Briefly, a synthetic peptide corresponding to rat VGF 556-564, with the addition of a C-terminal cysteine residue, was conjugated via its C-terminus to keyhole limpet hemocyanin (KLH), and used for immunizations.

### IHC and image analysis

Pancreas sections have been used to perform IHC. After a first treatment with Triton X-100 (0.1% in PBS 1× for 45 min.), the sections have been washed with PBS and incubated overnight with the guinea-pig TLQP antiserum diluted with PBS containing 30ml/L of normal donkey serum, 30ml/L of normal mouse serum and 0.02g/L NaN_3_. Guinea-pig/mouse insulin, rabbit glucagon (concession from Professors J.M. Polak and S.R. Bloom), rabbit somatostatin (AB_1143012), rabbit gC1q-R (AB_10675815) and C3a-R1 (AB_2687440) antibodies have been also used either single or in double staining (Table 1) while the relevant species-specific donkey secondary antibody/ies, conjugated with either Cy3, Cy2 or AMCA (anti guinea-pig AB_2340460; anti-mouse IgG, AB_2340460 /AB_2341099, anti rabbit: AB_2307443) were used to reveal immunoreactivity of the primary antibodies. Routine controls have been performed, included the substitution of each antibody/antiserum, in turn, with PBS or pre-immune sera and the testing of each secondary antibody with the respective non-relevant primary antibody/antiserum. Slides were covered-slipped with PBS-glycerol observed and photographed with BX51 fluorescence microscope (Olympus, Milan, Italy) equipped with Fuji S3 Pro digital Cameras (Fujifilm, Milan, Italy). For immunostaining quantification (through ImageJ software, US National Institutes of Health) of the TLQP staining, 6 animals per group have been selected and 3 photos (20x) of 3 different pancreatic islets (per each animal) clearly marked with the insulin antibody, have been taken in standard conditions (identical image acquisition settings and exposure times). The insulin immunstaining quantification has been similarly performed. The RGB/TIFF images have been first converted to 8-bit grayscale, then the boundaries of each islet have been manually traced by the user and, finally, the background fluorescent signal has been removed by a manual thresholding process. For each islet we have analyzed both the number of immunopositive cells and the image optical density.

**Table 1.** Antisera and antibodies used in the study.

| Antiserum/Ab | Producer | Species | Dilution |
| --- | --- | --- | --- |
| TLQPp | NEF-Laboratory, Cagliari, Italy | Guinea-pig | 1:1500 |
| Insulin | RPMS, London, UK; | Guinea-pig | 1:1500 |
| Insulin | RPMS, London, UK; | Mouse | 1:1000 |
| Glucagon | RPMS, London, UK; | Rabbit | 1:300 |
| Somatostatin | Abcam, Cambridge; UK | Rabbit | 1:300 |
| gC1q-receptor | Abcam, Cambridge; UK | Rabbit | 1:400 |
| C3a-receptor 1 | Abcam, Cambridge; UK | Rabbit | 1:50 |

Ab: antibody, TLQPp: TLQP peptides, NEF: Neuro-endocrine fluorescence, RPMS: Royal Postgraduate Medical School, Imperial College, gC1q: globular heads of C1q, C3a: complement component 3a

### ELISA

Competitive ELISA has been carried out as previously described in detail [8]. Briefly, multiwell plates (Nunc Thermo Scientific, Milan, Italy) have been coated with the relevant synthetic peptide and coating buffer (Na_2_CO_3_: 1.59 g/L, NaHCO_3_: 2.93 g/L, NaN_3_: 0.20 g/L; pH= 9.6) for 3 hours, hence treated with PBS containing normal donkey serum (90 ml/L), aprotinin (20 nmol/L), and EDTA (1 g/L) for 2 hours. Primary incubations (3h) have been carried out using guinea-pig TLQP antiserum (1:5k) followed by biotinylated secondary antibody (Jackson, West Grove, PA, USA, 1:10K,1h), streptavidin-peroxidase conjugate (Biospa, Milan, Italy, 30 min.), and tetramethylbenzidine substrate (X-traKem-En-Tec, Taastrup, Denmank, 100 ul/well). The reaction has been stopped with HCl (1 mol/L) and the optical density has been measured at 450nm using a multilabel plate reader (Chameleon: Hidex, Turku, Finland). Recovery of synthetic peptide (same used for immunization, plate coating and measurement standard) added to plasma, or to tissue samples at extraction was >85%. TLQP assay has been previously characterized [22] using multiple synthetic peptides, with IC_50_(pmol/ml) = 1.1; CV1=4% and CV2=6%.

### RP-HPLC analysis

Pooled extracts of additional control pancreas (n=6) and plasma (n=6) of the experimental groups were filtered with a 30 kDa cut-off Microcon filter (Merck Millipore, Tullagreen Carrigtwohill Co.Cork, Ireland) and dried in a Vacufuge Concentrator (Eppendorf, Milan, Italy), solubilized in 0.056% 2,2,2-trifluoroacetic acid (TFA) and spin at 14000 g for 5 minutes. As control, 0.5 nmol of rat/mouse TLQP-21 and TLQP-62 synthetic peptides (custom produced by CPC Scientific, Sunnyvale, CA, USA), were analysed by RP-HPLC (Dionex Ultimate 3000 instrument, Thermo Fisher Scientific, Sunnyvale, CA) and the extracted ion current peaks revealed by searching in the chromatographic profile their specific multiple-charged ions. The chromatographic column was a reversed phase Vydac (Hesperia, CA) C18 column with 5 μm particle diameter (250 × 10 mm). The following eluents were used: (eluent A) 0.056% (v/v) aqueous TFA and (eluent B) 0.05% (v/v) TFA in acetonitrile-water 80/20. The gradient applied was linear from 0 to 80% of B in 32 min, and from 80% to 100% of B in 1 min, at a flow rate of 0.20 ml/min. Tissue samples (50 μg) were similarly analyzed by RP-HPLC. The fractions eluted in the 6-36 minutes range, collected every two minutes from the outlet of the diode array detector, were dried with a Vacufuge Concetrator, and redissolved in PBS. Samples fractions eluting at the same position of the synthetic peptides (TLQP-21 and TLQP-62 standards) were analyzed by ELISA (for statistic evaluation, triplicate measurements were analysed) [23]. The protein concentation of the analysed fractions was detemined by the bicinchoninic acid assay (BCA assay, Thermo Scientific).

### Statistical analysis

For each experimental set, the normality of data distributions was preliminary checked using the Goodness-of-fit test. Resulting p values were >0.05 in all cases, hence the following parametric tests were appliedStatistical analysis has been performed using ANOVA, followed by a *post*-*hoc* test (Student-Newman-Keul), or paired two tailed Student’s T-test, through Microsoft Excel or SatistiXL softwares. When the *p* value was lower than 0.05, the considered groups have been evaluated significantly different from each other.

## RESULTS

### Pancreas

In normal endocrine pancreas (Fig. 1), TLQP antiserum stains peripheral (Fig. 1a,d) and central islet cells (Fig. 1g). The positive peripheral cells are identified as somatostatin (Fig. 1b,c) and glucagon (Fig. 1e,f) secreting cells. Instead, the internal TLQP positive cells contain, as expected, insulin (Fig. 1h,i). The insulin immunoreactivity is strong in control pancreas (Fig. 2a) but it is reduced, as predictable, at 30 (Fig. 2b) and 120 (Fig. 2c) min. after the glucose load, in agreement to the decrease in image optical density (*p*<0.001 and *p*<0.05; controls *vs.* 30 and *vs.* 120 min. respectively; Fig. 2g) and in number of cells/area (*p*<0.05; controls *vs.* 30 and *vs.* 120 min.; Fig. 2h). Likewise, TLQP immunoreactivity is well-visible in controls (Fig. 2d) but it disappears at 30 min. after the glucose load (Fig. 2e), returning to be observable at 120 min. (Fig. 2f). These results are in agreement with the decrease at 30 min. in image optical density (Fig. 2i: controls *vs.* 30 min.; *p*<0.05) and number of cells/area (Fig. 2j: controls *vs.* 30 min.; *p*<0.05). At 120 min. after the glucose load, the image optical density (Fig. 2i: 30 *vs.* 120 min.; *p*<0.05) and the number of cells/area (Fig. 2j: 30 *vs.* 120 min.; *p*<0.05) return similar to control levels. We have also studied the expression of the two TLQP-21 receptor molecules in the pancreas of both normal and glucose-treated mice. In the normal pancreas, labeling with gC1q-R is mainly found in peripheral cells of the islets (Fig. 3a) largely colocalized with TLQPp (Fig. 3b,c). Instead, the C3a-R1 staining is well represented in the entire islet, even if more represented in the central cells (Fig. 3d), hence only partially colocalized with TLQPp (Fig. 3e,f). The staining of the two receptors seems comparable between normal and glucose-treated animals (not shown). Raw data are included in S1_Fig.tif.

**Fig 1.**
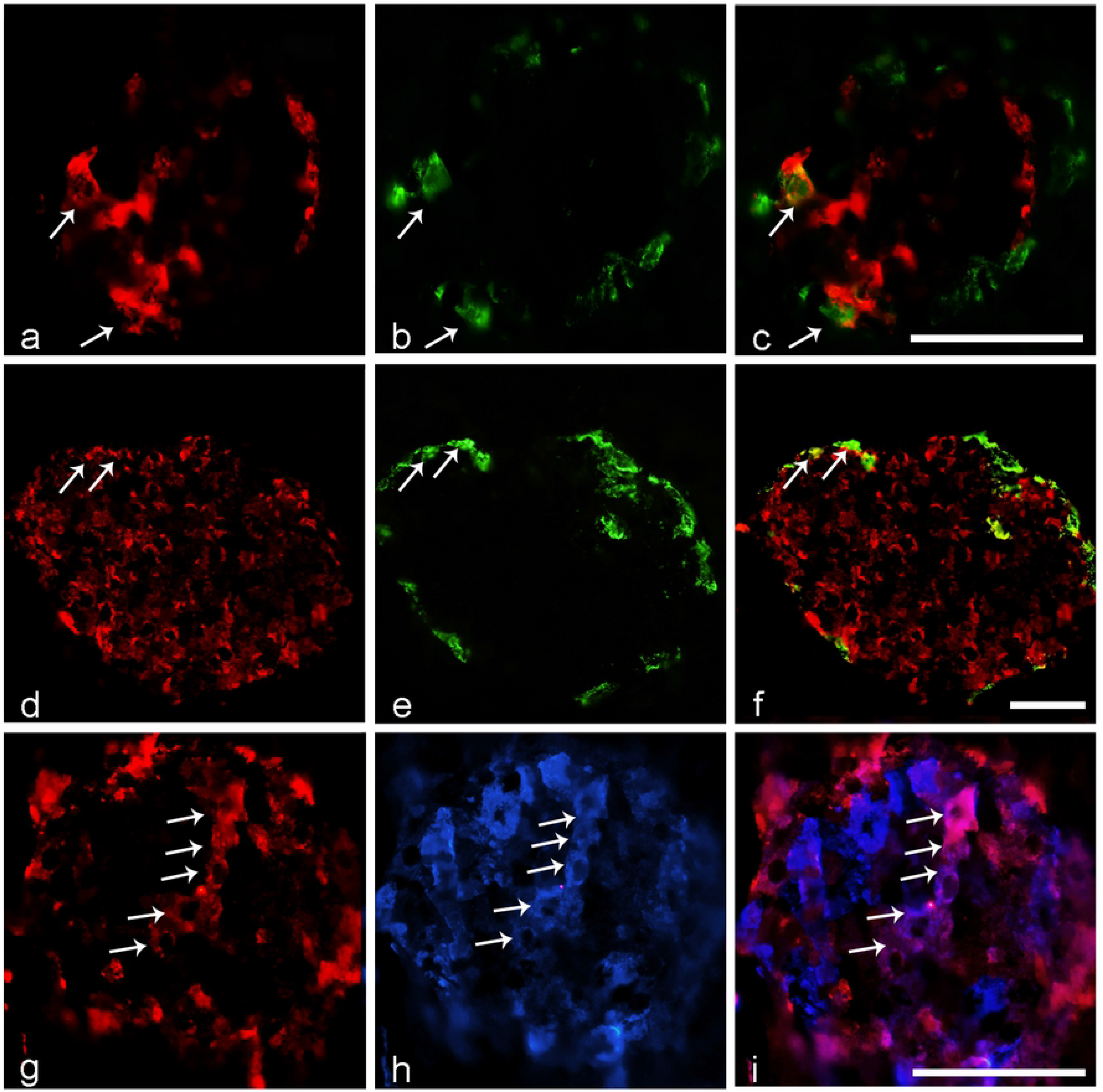
TLQP immunoreactivity in normal pancreas. The TLQP antiserum stains peripheral and central islet cells (**a**,**d**,**g**; Cy3 red labeling). Some of the peripheral cells are identified as somatostatin secreting cells (**b**: somatostatin antibody, Cy2 green labeling) as shown by double labeling (**c**: merge cells are identified by the arrows), some others contain glucagon (**e**: glucagon antibody: Alexa 488 green labeling) as confirmed by double staining (**f**; colocalized cells are identified by the arrows). The TLQP positive cells disposed in the centre of the islet are instead identified as insulin secreting cells (**h**: insulin antibody, AMCA, blue labeling) as confirmed by double staining (**i**: violet merged cells are identified by the arrows). Scale bars: 100um.

**Fig 2.**
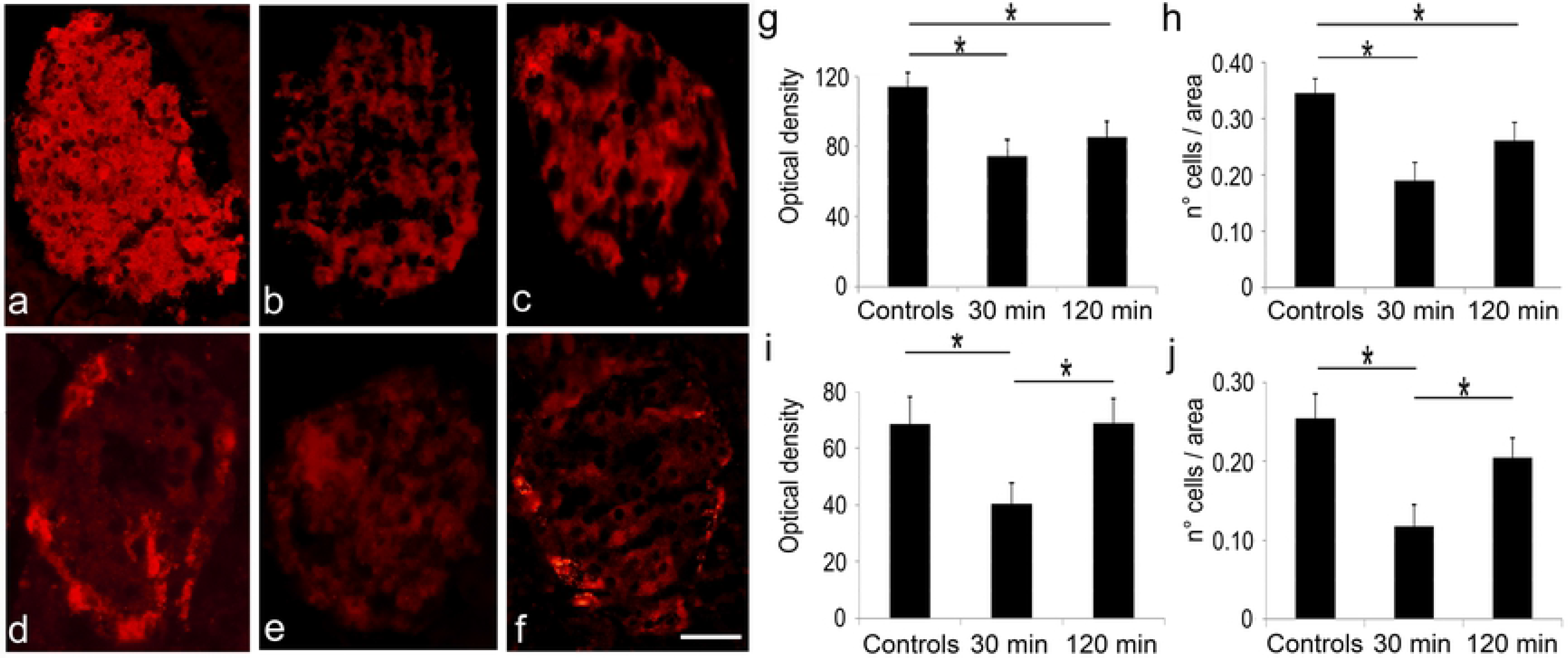
TLQP and insulin immunostaining in pancreas upon glucose load. As expected, the insulin antibody reveals the stronger staining in control mouse (**a**), compared to the weaker immunoreactivity observed upon glucose load, at both 30 (**b**) and 120 min. (**c**). TLQP antiserum labels bright cells in control mouse (**d**), but lost its immunoreactivity at 30 min. after a glucose load (**e**), returning to be visible at 120 min. (**f**). Insulin and TLQP antibodies: Cy3 red labelling. Scale bars: 40um. Immunostaining quantification using the insulin antibody shows that the image optical density (**g**) and the number (n°) of cells/area (**h**) are both reduced at 30 min. (*p*=0.001 and *p*=0.005, respectively), and also at 120 min. (*p*=0.033 and *p*=0.049, respectively). Immunostaining quantification using the TLQP antiserum reveals that the image optical density (**i**) and the number of stained cells (**j**) are both reduced at 30 min. (*vs.* control: *p*=0.024 and *p*=0.003, respectively) and re-increased at 120 min. (*vs.* 30 min.: *p*=0.027 and *p*=0.025, respectively).

**Fig 3.**
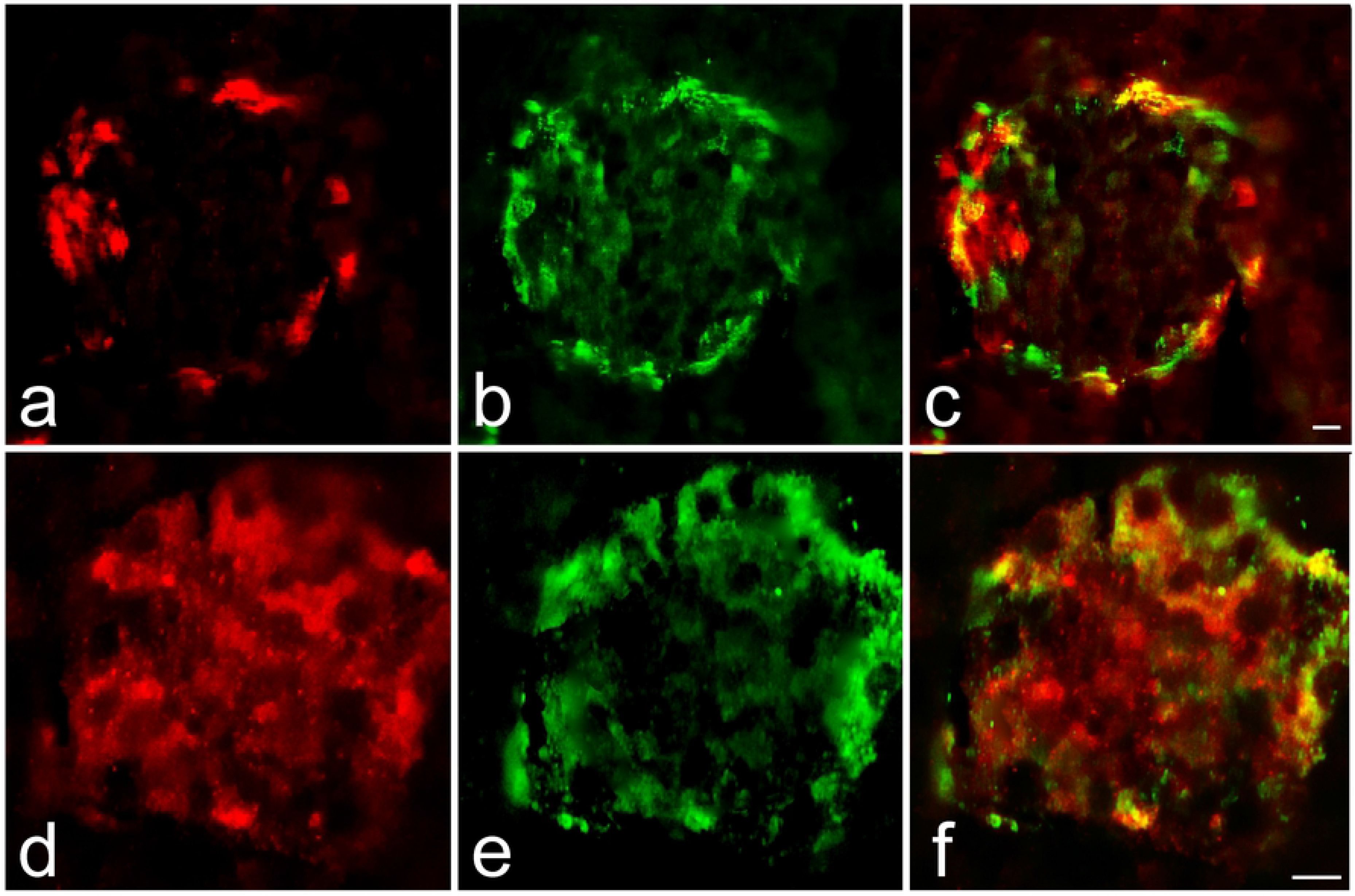
TLQP-21 receptors in pancreas. The gC1q-R (**a**: Cy3 red labeling) is expressed within the majority of peripheral TLQP positive cells (**b**: Cy2 green labeling) as confirmed by double labeling (**c**). Instead, the C3a-R1 (**d**: Cy3 red labeling) is found in peripheral as well as in a number of central cells, hence it is largely colocalized with TLQPp (**e**: Cy2 green labeling), as shown by double labeling (**f**). Scale bars: 20um.

### Plasma

As expected, after glucose load the glycaemia rises at 30 min. (Fig. 4a: controls *vs.* 30 min., *p*<0.001) to fall back at 120 min. (Fig. 4a: controls *vs.* 120 min. *p*<0.001; 30 *vs.* 120 min. *p*<0.001). Similarly, TLQP immunoreactivity increases at 30 min. after the glucose load (Fig. 4b: controls *vs.* 30 min., *p*<0.05), but reaches higher levels at 120 min. (Fig. 4b: controls *vs.* 120 min. *p*<0.001; 30 *vs.* 120 min.; *p*<0.05). Raw data are included in S1_ Fig.tif.

**Fig 4.**
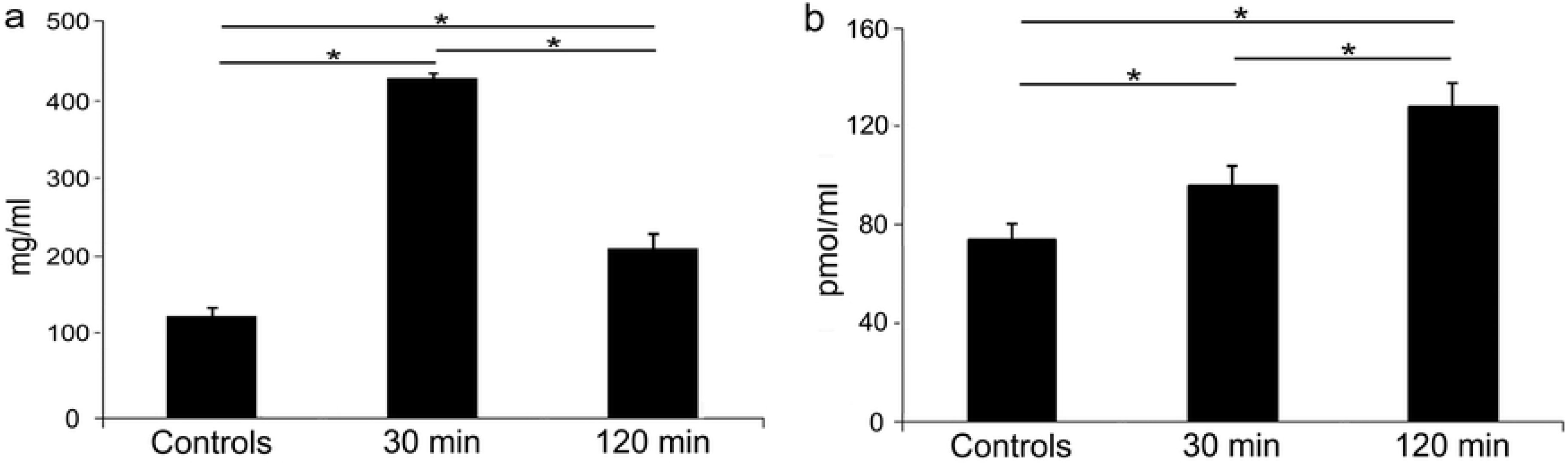
Glycaemia and TLQP immunoreactivty in plasma upon glucose load. Glycaemia (**a**) rises at 30 min. (*vs.* controls, *p*=0.001) to fall back after 120 min. (controls vs. 120 min. *p*=0.001; 30 vs. 120 min., *p*=0.001). Similarly, TLQP levels (**b**) ranging about 70–80 pmol/ml in normal conditions, increase at 30 min. (*vs.* controls: *p*=0.041), reaching higher values at 120 min. (controls vs. 120: *p*=0.001; 30 vs. 120 min.: *p*=0.049), mg/ml:milligrams/milliliters; pmol/ml:picomoles/milliliters.

### Reverse-phase HPLC

Upon HPLC using pooled pancreas extracts from the control group (Fig. 5), TLQP immunoreactivity is found by ELISA in a fraction at elution position coinciding with the synthetic TLQP-21. Using HPLC with plasma samples from the experimental groups (Fig. 6), TLQP immunoreactivity was always found by ELISA in fractions at elution positions coinciding with synthetic TLQP-21. After glucose load, TLQP immunoreactivity starts to increase in the fractions of the samples taken at 30 min, with a higher statistically significant peak in the fractions of the samples taken at 120 min (*vs.* controls p<0.05, Fig. 6 e), in agreement to the changes we revealed measuring the totality of the TLQP peptides (shown above). To strengthen this result, we also analysed pancreas (S2_Fig.tif) and plasma (S3_Fig.tif) fractions at elution positions coinciding with the extended TLQP-21 form, namely TLQP-62. However, we have not found any statistically significant changes between plasma fractions of control and treated samples (nor at 30 or at 120 min). The Fig. 7 resumes the TLQP-21 variations.

**Fig 5.**
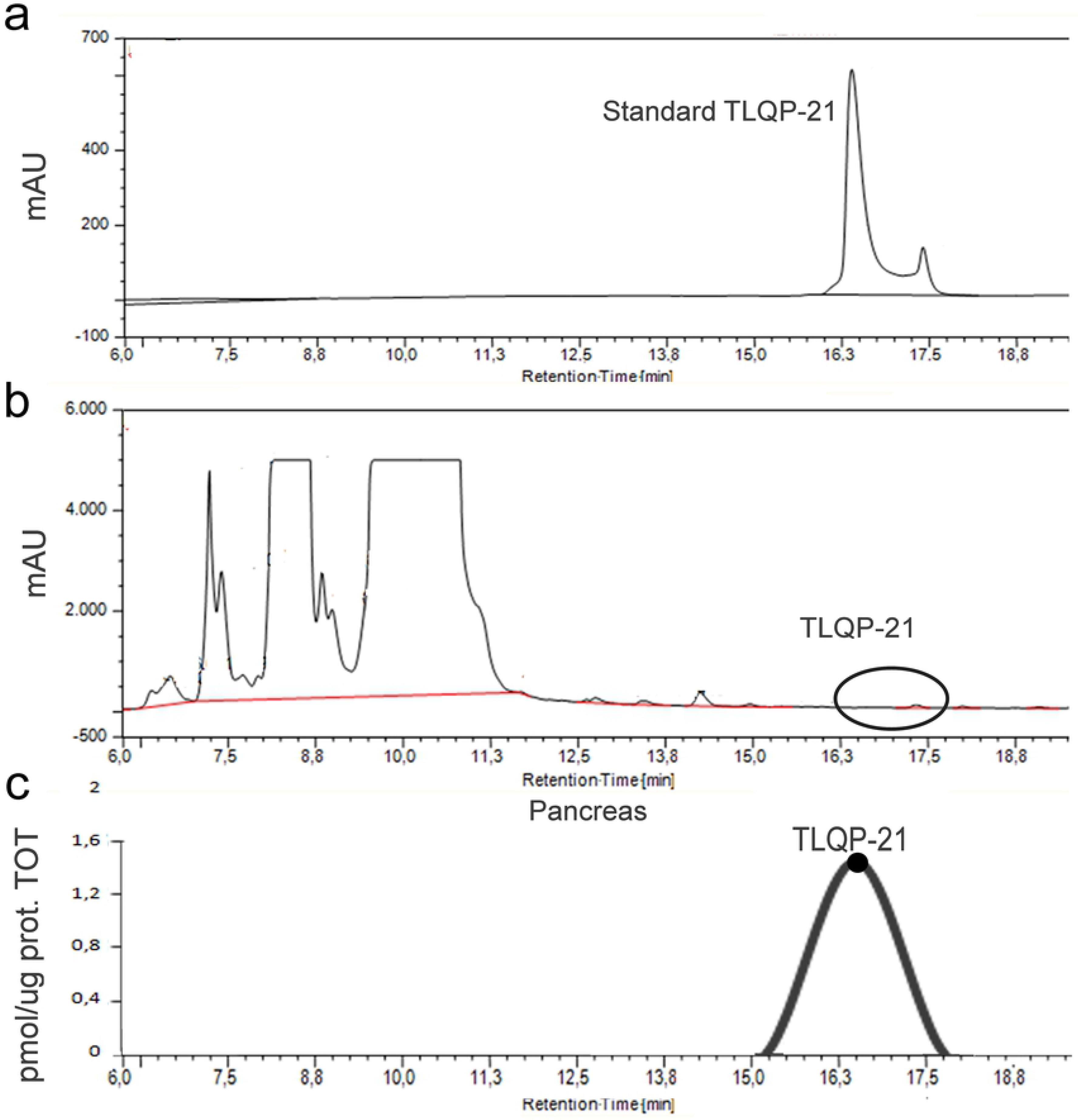
RP-HPLC coupled to ELISA (pancreas). Elution profile of TLQP-21 standard peptide (retention time 15-18 min) (a) and control pancreas (b) with evidenced the elution interval of fractions reactive with TLQP-antiserum by ELISA (c). pmol/ug prot.TOT: picomoles/micrograms of total proteins; mAU: milli-absorbance unit.

**Fig 6.**
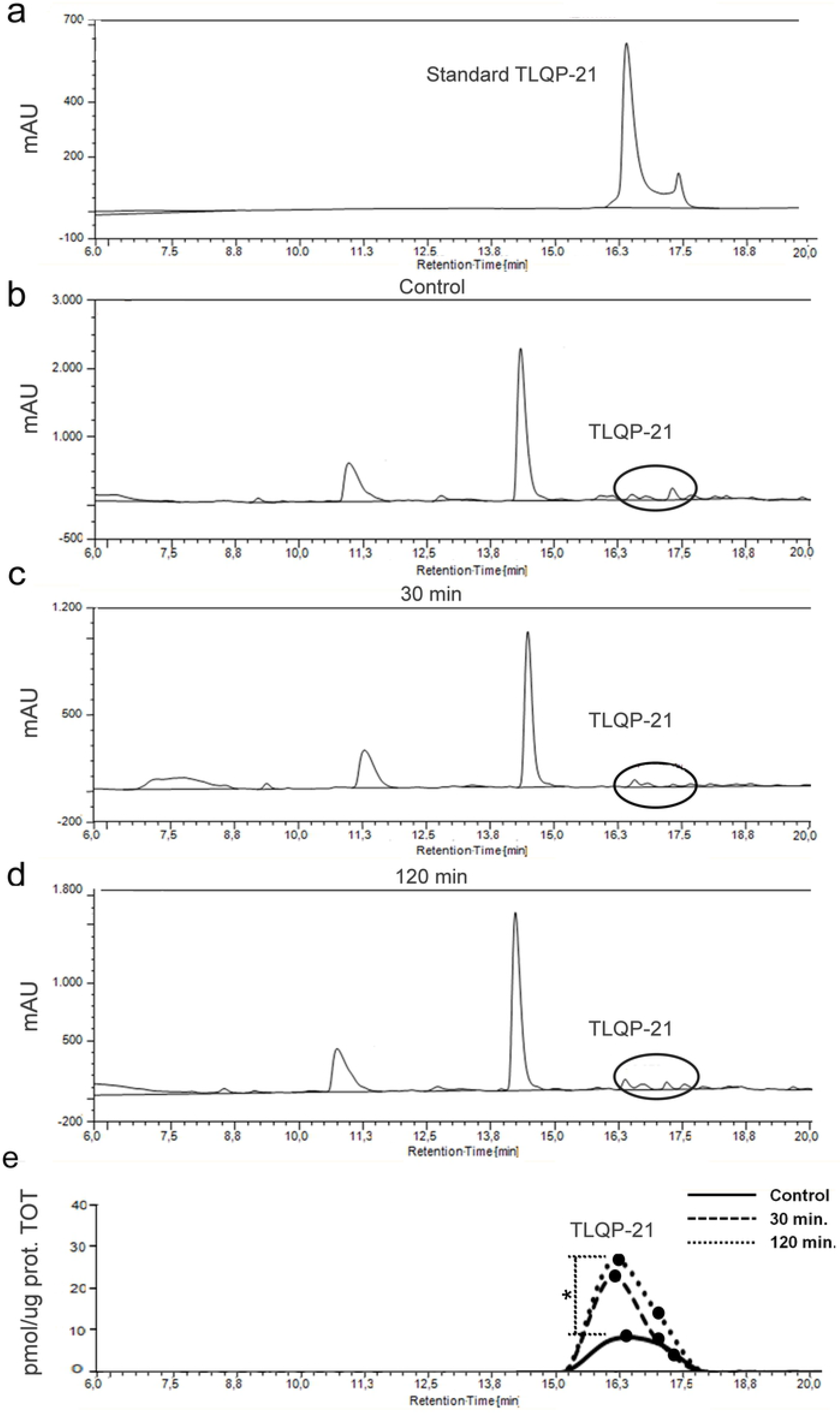
RP-HPLC coupled to ELISA (plasma). Elution profile of TLQP-21 standard peptide (retention time 15-18 min) (a), plasma samples in normal conditions (b) after 30 min (c) and after 120 min (d) of glucose load with evidenced the elution interval of fractions reactive with TLQP-antiserum by ELISA (e). TLQP-21 levels in plasma samples starts to increase at 30 min, reaching a statistically significant value at 120 min (e: *vs.* controls p<0.05). Pmol/ug prot.TOT: picomoles/micrograms of total proteins; mAU: milli-absorbance unit.

**Fig 7.** TLQP-21 changes. TLQP-21 variations are shown in parallel with glycaemic and insulin changes.

## DISCUSSION

At 30 min. after the glucose load, when the glycaemic peak is reached, the immunoreactivity of the entire population of TLQPp is scarcely detectable in the pancreas, concurrent with a TLQP-21 increase in plasma reaching a high peak at 120 min. Our results are in agreement with the role of TLQP-21 as hypoglycemic agent [11]. Indeed, TLQP-21 peripherally injected into rats, is able to reduce the blood-glucose peak (after about 20 min. from the bolus ingestion) and to increase the plasma insulin levels [11]. Obviously, TLQP-21 peptide in plasma can also originate from different organs/tissues. In fact, in addition to pancreatic islets [8, 9], TLQP-21 can also be released by the adrenal gland [19], stomach [20], pituitary-ovary axis [24], as well as (albeit less expected) median eminence [24]. Nonetheless, after a glucose load, the massive pancreatic degranulation of TLQPp at 30 min. in parallel with the starting of the TLQP-21 increase in plasma, should be in favor of considering the endocrine pancreas as the origin of at least part of the TLQP-21 amount in plasma. At 120 min., although the TLQP-21 return to be immunoreactive in the pancreas, it still increased in the plasma, as previously reported [9]. When plasma stability was measured, it was reported that TLQP-21 has a terminal half-life of about 110 min, hence roughly comprising our temporal range of measurements [25]. The major plasmatic increase of the TLQP-21 at 120 min rather than at 30min at the glycaemic peak is not clear, however we may hypothesize a modulatory activity of TLQP-21 at a late-phase of glucose-induced insulin secretion. The restore of the production in the pancreas could be compatible with a hypothetical re-accumulation phase, correlated to the *de novo* biosynthesis of TLQP-21. We have also found the presence of the two TLQP-21 receptors (gC1q-R and C3a-R1) in pancreatic cells. While limited information is so far available regarding the pancreas immunolocalization of gC1q-R [26], C3aR1 is known to be expressed in human and mouse islets, within β- and α- cells [27] and to potentiate glucose-induced insulin secretion from human and mouse islets [27]. Furthermore, C3a-R1, compared to gC1q-R, has been more studied as TLQP-21 receptor [13, 28, 29] and has been characterized in its precise reactivity depending on the C-terminal sequence of the TLQP-21 peptide [13]. We can hypothesize that TLQP-21, by returning to the pancreas from the plasma, could interact with the C3a-R1, in order to modulate the insulin secretion. Other hormones are known to act in this way, for example the gastrin peptide released from the stomach into the bloodstream, returns to the stomach in order to stimulate secretion and motility. In agreement with our data, targeted deletion of VGF in mice [30, 31] decreases islet cell mass, and reduces circulating insulin levels. Moreover, patients with obesity and Type 2 Diabetes (T2D) showed an altered TLQP response compared to normal weight subjects, while the chronic injection of TLQP-21 in pre-diabetic rats preserves islet mass and slows T2D onset [11]. In conclusion, we can speculate that, after glucose stimuli, TLQP-21 could modulate the insulin secretion through an endocrine activity.

## Supporting informations

**S1 Fig. Raw data.** Pancreas image analysis (optical density and number of cells/area) and ELISA analysis of plasma samples.

**S2 Fig. RP-HPLC of pancreas coupled to ELISA.**
Elution profile of TLQP-62 standard peptide (retention time 22-25 min) (**a**), and control samples (**b**) with evidenced the elution interval of fractions reactive with TLQP-antiserum by ELISA (**c**). Pmol/ug prot.TOT: picomoles/micrograms of total proteins; mAU: milli-absorbance unit.

**S3 Fig. RP-HPLC of plasma coupled to ELISA.**
Elution profile of TLQP-62 standard peptide (retention time 22-25 min) (**a**), samples in normal conditions (**b**) after 30 min (**c**) and after 120 min (**d**) of glucose load with evidenced the elution interval of fractions reactive with TLQP-antiserum by ELISA (**e**). The increase of TLQP-62 at 30 and 120 min is not statistically significant (**e**: 30, 120 min *vs.* controls p>0.05). Pmol/ug prot.TOT: picomoles/micrograms of total proteins; mAU: milli-absorbance unit.

## ACKNOWLEDGMENTS

This work was supported by grants of the University of Cagliari (FIR, Cocco), Professors J.M. Polak and S.R. Bloom are thanked for antibodies.

## Author Contribution

GC, CC and G-LF conceived and designed the experiments; GC, BN, CB, BM, FD, MP performed experiments; GC, CC analyzed the data; GC and CC wrote the paper; FN, BM contributed reagents/materials/analysis tools.

